# Small-scale, diverse horticultural systems and semi-natural grasslands support complementary pollinator populations

**DOI:** 10.1101/2025.02.27.640499

**Authors:** Jutta Crois, Kris Verheyen, Ivan Meeus, Maxime Eeraerts

**Affiliations:** Forest & Nature Lab, Department of Environment, Faculty of Bioscience Engineering, Ghent University, Geraardsbergse steenweg 267, Melle-Gontrode, Belgium; Laboratory of Agrozoology, Department of Plants and Crops, Faculty of Bioscience Engineering, Ghent University, Coupure links 653, Gent, Belgium; Aculea, Hymenoptera study group, Natuurpunt, Coxiestraat 11, Mechelen, Belgium

## Abstract

Pollinating insects are of great importance in natural ecosystems and agricultural landscapes. Despite their crucial role for food production, safeguarding pollination services is becoming increasingly challenging due to pollinator declines. Habitat loss and agricultural intensification are among the main drivers, causing a substantial reduction in the availability of floral resources. Despite efforts, the uptake of measures to halt these declines in agricultural landscapes is low. Implementation of a novel crop production system is therefore proposed as a solution to ensure food security whilst concomitantly minimising environmental degradation and the associated biodiversity loss. Agricultural diversification is one such strategy that is known to promote multiple ecosystem services. Yet, the effects of agricultural diversification on pollinator populations in farm-scale, observational studies remains understudied.

Here, we examine the importance of small-scale, diverse horticultural systems (SDHS) to support wild pollinator communities. Floral resources and wild pollinators were sampled in both SDHSs and semi-natural grasslands (SNG) in 16 landscapes in Flanders, Belgium. During late summer, these SNGs provide and important source of floral resources, thereby serving as a benchmark habitat to which the SDHSs are compared.

Our study highlights the value of SDHSs as they provide diverse and abundant floral resources compared to SNGs. Moreover, pollinator species richness and abundance were comparable between the two habitats. Conversely, floral resource and pollinator community composition differed, indicating that different pollinator communities used the complementary resources provided by both habitat types. Furthermore, we show that SNGs represent the most specialised habitat type. We therefore argue that SDHSs can be considered to be an additional, complementary source of floral resources to those provided by SNGs. However, these SDHSs cannot substitute SNGs, given the importance of these grasslands for pollinators.

## Introduction

More than 80% of the world’s flowering angiosperms and 75% of all crops rely to some extent on animal-mediated pollination (Klein et al., 2007; Ollerton et al., 2011). Besides, the global area of pollinator-dependent crops has increased rapidly in recent years (Aizen et al., 2019). Therefore, both managed and wild pollinating insects represent a crucial component in agriculture. The average contribution of the Western honeybee (*Apis mellifera*), the most commonly managed pollinator, to crop production is estimated to be $3000 per hectare (Kleijn et al., 2015; Osterman et al., 2021). Nevertheless, the efficacy of crop pollination by honeybees does not always reach the same levels as those observed for wild insects, for which the relationship with crop pollination is consistently positive (Garibaldi et al., 2013). Hence, wild pollinators are considered at least as important for crop production and yield as honeybees (Eeraerts, DeVetter, et al., 2023; Reilly et al., 2024).

Safeguarding pollination services is becoming increasingly challenging due to declines of wild pollinator populations (Hallmann et al., 2017; Seibold et al., 2019). Multiple factors underlie this insect decline and these factors often interact (Goulson et al., 2015). Among the main drivers are habitat loss and agricultural expansion and intensification (Kleijn et al., 2009; Seibold et al., 2019). Since the 1940s, agriculture has undergone significant expansion and intensification: farms declined in number but expanded in size, pesticides and inorganic fertilizers were increasingly applied, monocultures of the same crop for several consecutive years replaced crop rotations, and hedgerows and other semi-natural habitat (SNH) elements were largely removed from the agricultural landscape (Robinson & Sutherland, 2002). Consequently, this agricultural intensification caused a considerable reduction in plant abundance and diversity in agricultural landscapes, thereby reducing the available floral resources for insects (Carvell et al., 2006; Scheper et al., 2014). Hence, both wild insects and the ecosystem services they provide (e.g. pollination, pest control), are negatively affected in intensive agricultural landscapes (Dainese et al., 2019; Eeraerts et al., 2017).

In order to provide adequate support for pollinator populations and the services they deliver, the amount of SNH in the surrounding landscape should be sufficiently high. It is recommended that the proportion of SNH in agricultural landscapes should reach a minimum of 15% (Eeraerts, 2023). Beyond this threshold, the quality of SNH, rather than quantity, will have a greater impact (Fijen et al., 2024). SNHs provide nesting sites and floral resources over space and time, thereby maintaining rich pollinator communities (Martínez-Núñez et al., 2022). Indeed, different habitats vary in their relative importance to pollinators during their active period: woody vegetation (e.g. forest edges) highly contributes to the overall landscape-level floral resource abundance in early spring, while herbaceous habitats (e.g. grasslands) gain importance later in the season (Ammann et al., 2024; Maurer et al., 2022). Without this phenological continuity, a discrepancy between floral resource provision and pollinator needs throughout their active period would emerge (Mandelik et al., 2012; Timberlake et al., 2019).

Previous research has shown that the agricultural matrix can complement SNHs. Woody SNHs and fruit orchards, for instance, are characterized by a distinct set of nesting and floral resources, thereby complementing each other during the flight season of pollinators (Eeraerts, Van Den Berge, et al., 2021). Cultivating a variety of mass-flowering fruit crops, each having distinct, sequential flowering phenologies, also improves floral resource provision for pollinators (Martins et al., 2018). However, increasing crop compositional heterogeneity does not always increase the abundance of pollinators, and can even negatively affect their numbers (Hass et al., 2018; Pisman et al., 2022).

At a local scale, pollinators can be enhanced by sowing (wild)flower strips or planting hedgerows, as these linear elements produce high amounts of nectar per unit area (Timberlake et al., 2019; von Königslöw et al., 2022). Experimental research also showed that the frequency of pollinator visits to crops increased with the presence of flower strips (Feltham et al., 2015). However, the uptake of these and other habitat conservation or creation measures by farmers remains relatively limited (Kleijn et al., 2019). The implementation of pollination-enhancing measures such as flower strips and hedgerows is often constrained by various factors, including space requirements, financial commitments and time availability (Eeraerts et al., 2020). Indeed, although these agri-environmental schemes have been shown to increase biodiversity, land has to be taken out of production (Batáry et al., 2015). Hence, the ecological value of management practices supporting biodiversity, and pollinators in particular, often does not outweigh the economic uncertainty (Scheper et al., 2023).

A promising strategy for pollinator conservation in agricultural landscapes is crop diversification, a measure by which the availability of floral resources increases both in space and time, without taking land out of production (Fijen et al., 2025). Many different practices can be regarded as crop diversification measures. The sequential flowering of different cultivars, for instance, has been observed to increase pollinator abundance and species richness (Chabert et al., 2024; Eeraerts, 2022). Integrating shrubs and trees, a practice known as agroforestry, has also been found to enhance pollinator populations (Kay et al., 2020). Moreover, different foraging habitats can be provided by intercropping with herbaceous crops (Guariento et al., 2024; Kirsch et al., 2023; Pereira et al., 2015). Nevertheless, despite its potential value for pollinator management, there is only little research available on the effects of crop diversification on pollinators and the services they deliver (Fijen et al., 2025; Tamburini et al., 2020).

In this study, we want to investigate the role of small-scale, diverse horticultural systems (SDHS) for wild pollinating insects in late summer. Most of the farms that were visited, belong to the “CSA-netwerk” of Flanders and were situated in (peri-)urban areas. Previous research has shown that (peri-)urban landholdings are generally small-scaled, include a high diversity of crops, and can be highly productive, while simultaneously providing environmental and societal benefits. Yet, there is a general lack of information regarding the impact of these food producing systems on biodiversity and pollinator populations in particular (Nicholls et al., 2020). In late summer, when floral resource provision is low, semi-natural grasslands (SNG) are the main habitat contributing to the floral resource availability for pollinators. However, this provision often does not suffice (Timberlake et al., 2019). Therefore, we address the following research questions:

1. Can SDHSs complement SNGs in providing floral resources for wild pollinating insects in late summer
2. To what extent do SDHSs support communities of wild pollinators in late summer and how does this relate to SNGs?
3. Does a plant-pollinator network reveal complementary habitat use by wild pollinating insects in late summer?

## Materials and methods

### Study design and site selection

We selected a total of sixteen landscapes, in which a paired SDHS and SNG site were present, as habitat sites for data collection in Flanders, Belgium. In this region, intensive agriculture is widespread and forms an important economic activity (Jacquemin et al., 2017). The distance between the different landscapes ranged from 1.5 km to 109.2 km (Fig. 1)

**Figure 1:**
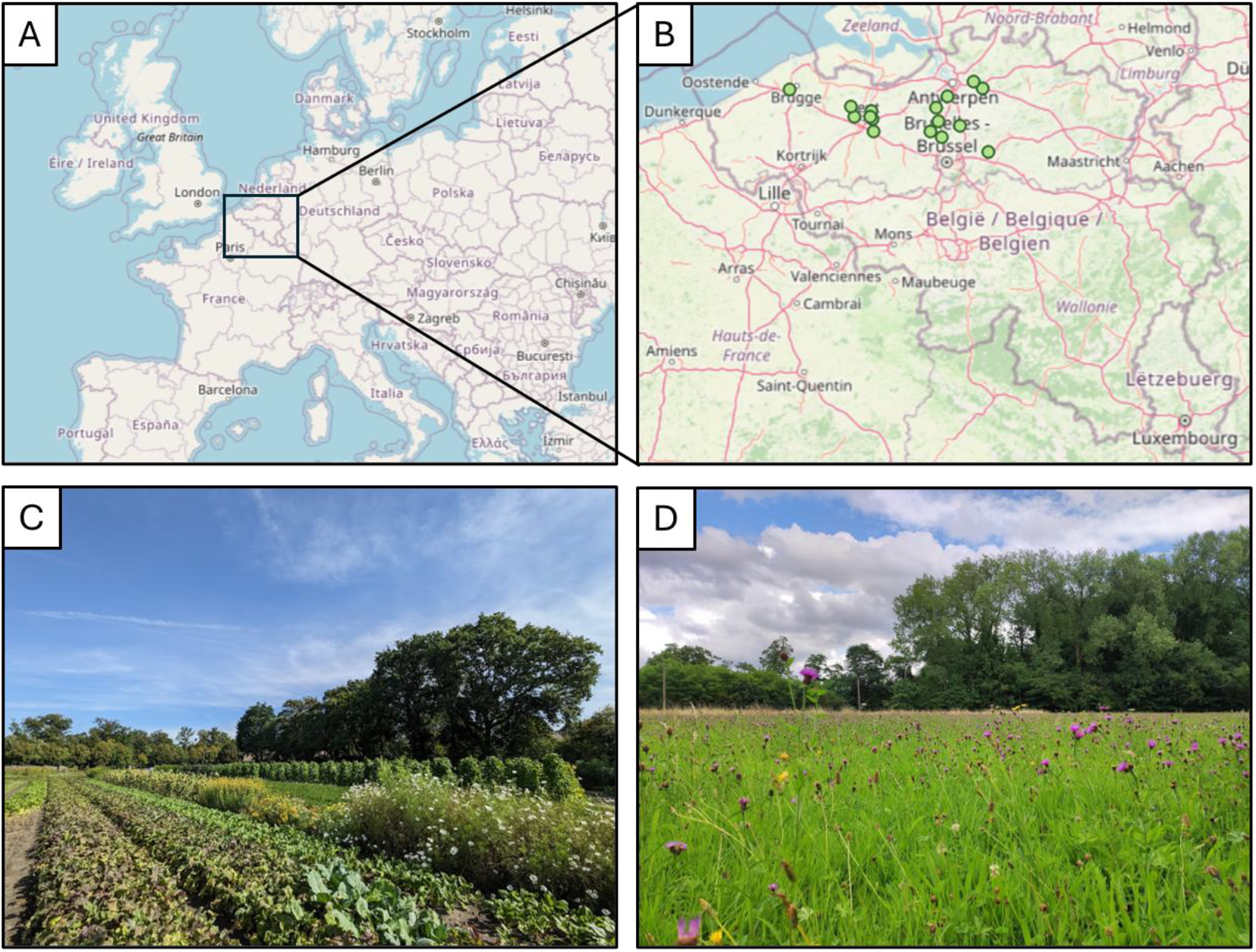
Overview of the study area and sampling habitats. Location of the study area in Europe (A) and an overview of the locations of the different paired habitat sites (n = 16) in Flanders, the northern region of Belgium (B). Example of a small-scale, diverse horticultural system (C) and semi-natural habitat (D).

In each landscape, a SDHS and SNG were chosen. SDHSs represent highly diverse, mixed food producing systems and most of the SDHSs we visited were part of the “CSA-netwerk” (Community Supported Agriculture Network). These farms are typically smaller than the average Flemish farms. They produce a diversity of fruits, vegetables, herbs and flowers, which are marketed through home sales, packets, the share-and pick principle, or a combination, and some of them also have livestock (Savels et al., 2024). Most of the SDHSs were certified organic farms, but usage of chemical pesticides and fertilizers is generally low or negligible on all these farms.

A high-value SNG was searched in a buffer zone with radius 1000 m around the SDHS. The SNG was selected based on the Biological Validation Map of Flanders, which classifies the vegetation type of nearly every parcel in Flanders and evaluates its biological value as “not valuable”, “biologically valuable” and “biologically very valuable” (De Saeger et al., 2023). All selected SNGs were biologically valuable and/or very valuable species-rich grasslands, mesophilic hay meadows and rough grasslands. In the final selection, the distance between the SDHS and SNG within each landscape ranged from 51 m to 1304 m (mean ± SE = 749 ± 83).

### Data collection

In each SDHS and SNG, two linear transects of 50 m^2^ (25 m x 2 m) were laid out for data collection. These transects were positioned so that they covered the representative vegetation of the whole parcel. Data of floral resources and flower-visiting pollinators were collected between the 25^th^ of July and 5^th^ of September 2023.

### Floral resources

Within each 50 m^2^ transect all plant specimen were identified to species level. If needed, we used the identification application ObsIdentify, provided that the observation certainty was at least 90% (Bishop et al., 2024). If species identification remained unknown, plant genus was noted and a plant specimen was taken to the lab for further identification. Five plant species remained unidentified and were named “Plant X” (with X = 1, 2, 3, 4 or 5). Only one of them was visited by pollinators (“Plant 1”; Table S1). Plant species in every transect were classified according to their phenology (i.e. vegetative, flowering, bloomed) and the number of flowerheads of all flowering plants was counted. These flower counts took into account the different types of inflorescences (e.g. single flowers for *Ranunculus acris*, umbels for *Achillea millefolium*).

### Pollinator survey

After the vegetation survey we collected data on the flower-visiting insects, hereafter referred to as “pollinators”. Along each transect, all pollinators, contacting the reproductive organs of a plant, were captured with an insect net for 40 minutes. To ensure adequate pollinator activity, surveys were conducted between 10:30 am and 18:00 pm, during dry and warm weather conditions (min. 17°C) with no or low wind (Table S2). We focussed on bees, hoverflies, butterflies and wasps. Sampling was carried out by JC and ME. Captured pollinators were kept in tubes during sampling and we kept track of the plant species on which they were foraging and caught. After each survey, all individuals were identified to species and the number of individuals per species captured on each flowering plant species was noted. If individuals could be identified to species in the field, they were released. Otherwise, they were taken to the lab for later identification. Identification was done using the identification key of Falk & Lewington (2017) for bees, Bot & Van de Meutter (2019) for hoverflies, and Wynhoff et al. (1999) for butterflies. Bee and hoverfly species identification was verified by JD and JH, respectively. All wasp specimens were identified by AD.

### Data analysis

#### Floral resources

Vegetation data of the two transect were pooled per SDHS and per SNG for analyses. The number of unique plant species per habitat type and the total number of flowerheads for each plant species were used as measure of species richness and flower abundance, respectively. We restricted the analysis of vegetation data to flowering and visited plant species, as we are interested in floral resource provision for pollinators. This decision is supported by the significant correlation between species richness and flower abundance of the flowering plants and the flowering and visited plants (Spearman’s rank correlation: ρ = 0.71, S = 1603, p < 0.001 and ρ = 0.99, S = 58, p < 0.001 for floral species richness and abundance, respectively).

To test the effect of habitat type on species richness, we used linear mixed-effect models (LME; function *lme*, R package *nlme*, Pinheiro et al. (2025)). A generalized linear mixed-effect model (GLMM; function *glmer*, R package *lme4*, Bates et al. (2025)) with Poisson distribution was used to investigate the effect of habitat type on flower abundance. In both models, we used habitat type as explanatory variable and sampling location ID as random factor.

To determine the effect of habitat type on the vegetation composition, a community composition dissimilarity among pairs of every site per habitat type was computed, based on the flower abundance data. We chose the Morisita-Horn dissimilarity index, as this method better accounts for the presence of rare species (Beck et al., 2013). Based on these dissimilarities distance matrices, a permutational multivariate analysis of variance was applied to test if habitat type affected the vegetation community composition (PERMANOVA, Anderson (2001); function *adonis2*, R package *vegan*, Oksanen et al. (2025)). The number of iterations was set at 1000. To visualize the community composition per habitat type, we used nonmetric multi-dimensional scaling (NMDS; function *metaMDS*, R package *vegan*). Additionally, we determined indicator species for both habitat types and computed the point-biserial correlation coefficients (rr_pb_; function *multipatt*, R package *indicspecies*; De Cáceres et al. (2010))

#### Pollinators

To evaluate how different habitat types support wild pollinators, we considered the following response variables: species richness of all pollinators, species richness of bees, species richness of hoverflies, abundance of all pollinators, abundance of bees and abundance of hoverflies. Honeybees were excluded from the analyses, as the focus was on wild pollinators. Pollinator data of the two transects were pooled per SDHS and per SNG for analyses. We analysed the effect of habitat type on pollinator abundance and species richness with GLMMs with Poisson distribution (function *glmer*, R package *lme4*). Habitat type was used as explanatory variable and location ID as random factor.

The effect of habitat on the pollinator community composition was determined similarly to the floral resources approach above. To calculate the community composition dissimilarity among pairs of sites, we used the Morisita-Horn index based on the pollinator abundance data. We used the dissimilarities distance matrices obtained to perform PERMANOVA to test if there is an effect of habitat type on the pollinator community composition (function *adonis2*, R package *vegan*). The number of iterations was set at 1000. Next, multivariate homogeneity of groups dispersion was calculated. The results were visualized in an NMDS plot (function *metaMDS*, R package *vegan*). Additionally, we determined indicator species per habitat type (function *multipatt*, R package *indicspecies*).

#### Plant-pollinator network

To assess whether SDHSs are complementary habitats to SNGs, we combined the plant and pollinator data in a plant-pollinator network and examined the degree of specialisation of these networks. Plant-pollinator networks were converted to network matrices (function *frame2webs*, package *bipartite*, Dormann et al. (2024)) which were then used to calculate the Blüthgen Hr_2_’ (function *networklevel*, package *bipartite*). Blüthgen Hr_2_’ is a network-level index that estimates the degree of specialisation of an entire network. The value of this index ranges between 0 and 1, with a value of 0 representing extreme generalisation and a value of 1 representing extreme specialisation (Blüthgen et al., 2006). Here, each combination of a sampling location and habitat type served as a single network, resulting in a total of 32 networks for which the Hr_2_’ was calculated. Subsequently, we allocated the Hr_2_’ values of each network to their corresponding habitat type, resulting in sixteen networks for each of these habitat types. The effect of habitat type on the degree of specialisation was evaluated using a LME (function *lme*, R package *nlme*), with Hr_2_’ as the response variable, habitat type as the explanatory variable and location ID as random factor.

#### Data assumptions and model validation

Outliers of response variables were identified by making a boxplot (function *boxplot*, package *graphics*, Murrell (2020)) and response variables were checked for a normal distribution by making a histogram (function *hist*, package *graphics*), dot chart (function *dotchart*, package *graphics*) and Q-Q plot (function *qqnorm, qqline*, package *stats*, Bolar (2019)) and by performing a Shapiro-Wilk normality test (function *Shapiro*.*test*, package *stats*). LMEs were chosen if data was normally distributed while GLMMs with Poisson distribution were used for count data. To validate the output of LMEs, we checked the residuals for outliers and normality as described above. For GLMMs we performed an additional model validation (function *testDispersion, simulateResiduals, testUniformity, testOutliers*, package *DHARMa*, Hartig (2022)). In case of outliers of the response variables, we tested the models with and without the outliers. All analyses were done in R version 4.4.1 (R Core Team, 2024).

## Results

### General

In total we identified 154 flowering plant species, of which 100 were visited at least once by a pollinator (Table S1). Of the visited flowering species, 51 were unique to the SDHS habitat, 33 were unique to the SNG habitat, and 16 species occurred in both habitats.

Overall, we caught 2138 pollinator specimens, of which 245 were honeybees. Regarding the 1893 wild pollinators, 945 were caught in the SDHSs and 948 in the SNGs. A total of 134 unique pollinator species were found, 85 species at the SDHSs and 98 at the SNGs. Among all these pollinators, 1038 are wild bees from 40 unique species, 658 are hoverflies from 44 species, 80 are butterflies from 11 species, and 117 are wasps from 39 species. Of the pollinator species, 36 are unique to the SDHS habitat, 49 are unique to the SNG habitat, and 49 species occur in both habitats.

### Flower and pollinator richness and abundance

We found that habitat type has no significant effect on the species richness of the flowering and visited vegetation, but has a significant effect on the flower abundance (Table 1, Fig. 2A, 2B; mean & SE, Table S3). There were some outliers in the flower abundance dataset, but removing these datapoints did not change the results. The species richness and abundance of all pollinators, bees and hoverflies are similar between the two habitat types (Table 1, Fig. 3A, 3B, 3D, 3E, 3G, 3H; mean & SE, Table S3). We detected outliers except for the hoverfly abundance dataset. However, the exclusion of these outliers did not change the results, and thus they were retained for further analysis (Table S5).

**Table 1:**
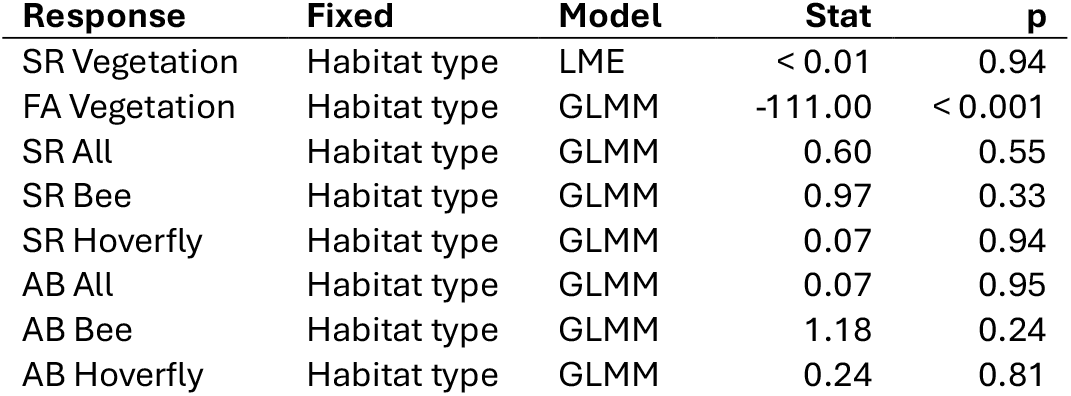
Summary of the results of the linear mixed-effect models (LME) and generalized linear mixed-effect models (GLMM) for the effect of the habitat type on the species richness (“SR”) and flower abundance (“FA”) of the flowering and visited plant species and on the species richness and abundance (“AB”) of all pollinators, bees and hoverflies. “Response” and “Fixed” are the response variable and fixed variable of the model, respectively. The reported statistics (“Stat”) are the F-value for LME and z-value for GLMM, together with the corresponding p-value (“p”) for each model. SDHS was used as reference habitat type in the models.

**Figure 2:**
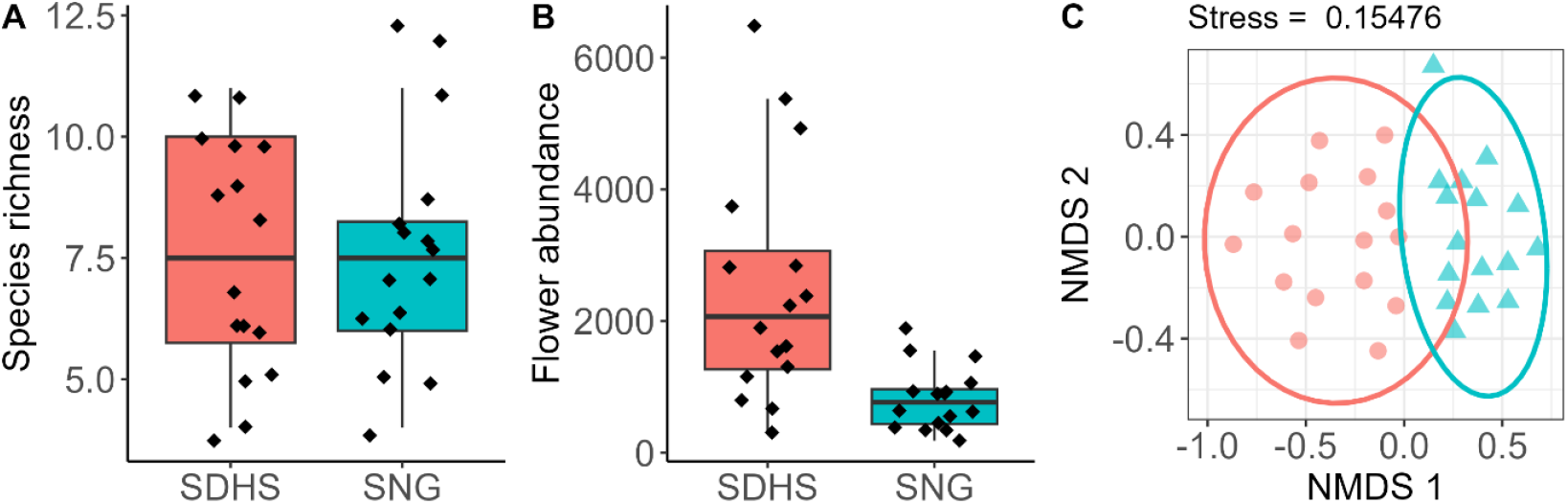
Boxplot of the species richness (A), boxplot of the flower abundance (B) and NMDS-plot of the flower abundance of the flowering and visited vegetation for the SDHS (pink) and SNG (blue) habitat type.

**Figure 3:**
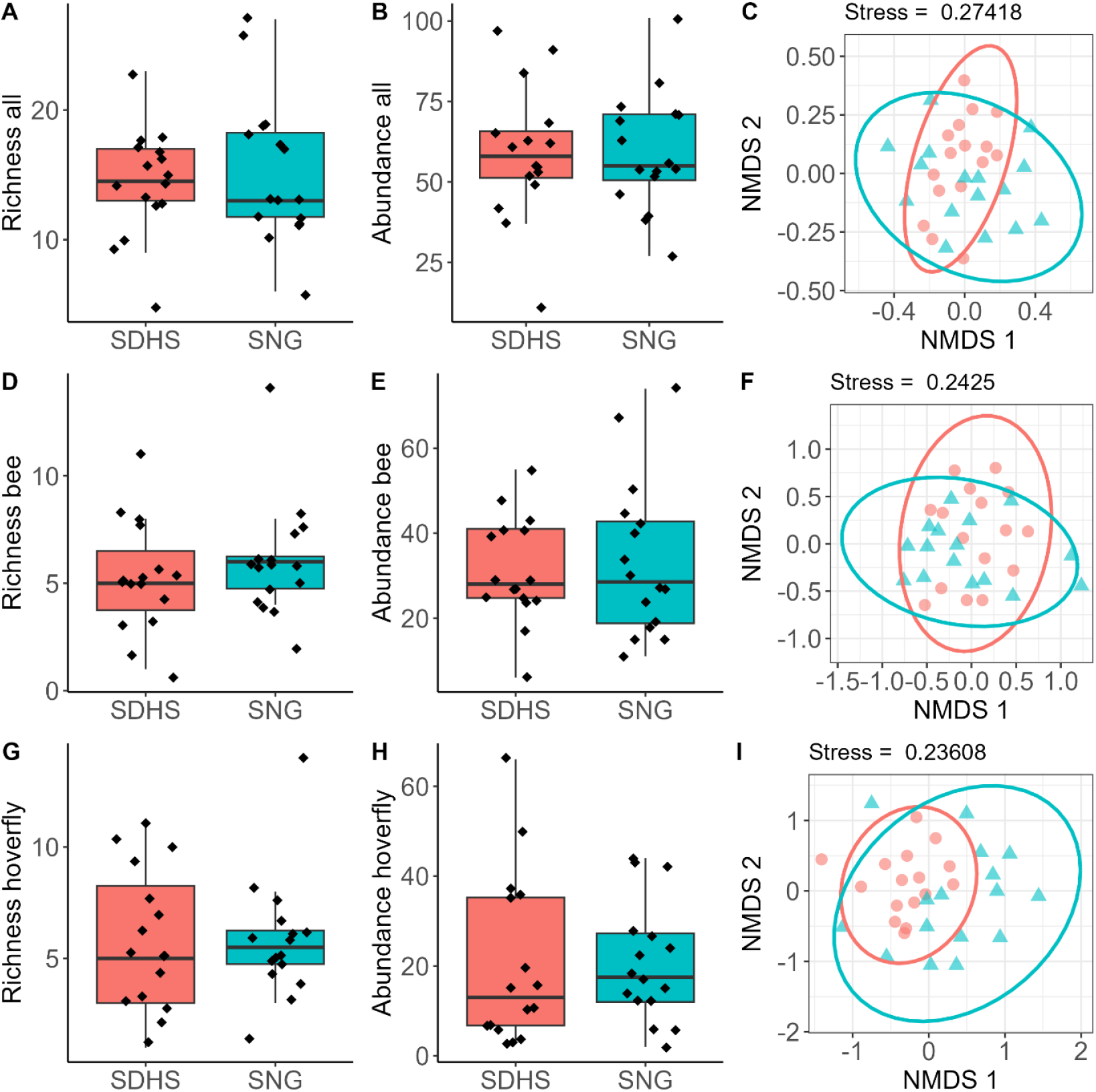
Boxplots of the species richness (left), boxplots of the abundance (middle) and NMDS-plots of the abundance (right) of all pollinators (A-C), bees (D-F) and hoverflies (G-I) for the SDHS (pink) and SNG (blue) habitat type.

### Vegetation and pollinator composition

Habitat type has a significant influence on the flowering and visited plant community composition (F = 3.39, p < 0.001; Fig. 2C). We identified *Galinsoga quadriradiata* (rr_pb_ = 0.33, p < 0.01) and *Phacelia tanacetifolia* (rr_pb_ = 0.27, p = 0.043) as indicator species for the SDHS habitat, and *Lotus corniculatus* (rr_pb_ = 0.45, p < 0.01), *Heracleum sphondylium* (rr_pb_ = 0.38, p = 0.019), *Lythrum salicaria* (rr_pb_ = 0.35, p < 0.01), *Cirsium arvense* (rr_pb_ = 0.35, p = 0.021), *Jacobaea vulgaris* (rr_pb_ = 0.31, p = 0.037 and *Centaurea jacea* (rr_pb_ = 0.28, p = 0.043) as indicator species for the SNG habitat.

Habitat type also clearly affects the community composition of bees (F = 2.30, p = 0.016; Fig. 3F) and hoverflies (F = 2.91, p = 0.024; Fig. 3I), but not the community composition of all pollinators (F = 1.98, p = 0.07; Fig. 3C). For the SDHSs we identified *Hylaeus communis* (rr_pb_ = 0.38, p = 0.046), *Syritta pipiens* (rr_pb_ = 0.38, p = 0.031) and *Pieris rapae* (rr_pb_ = 0.34, p = 0.039) as indicator species, while *Melitta nigricans* (rr_pb_ = 0.41, p < 0.01), *Helophilus pendulus* (rr_pb_ = 0.39, p = 0.032) and *Eristalis nemorum* (rr_pb_ = 0.36, p = 0.027) were indicator species for the SNGs.

Additionally, we calculated the mean values of flower abundance for each plant species and the mean values of visits for each pollinator species by summing the floral abundance or visits per species and dividing this value by 16 in order to take the absence of certain species at some locations into account. These mean values were then divided by the sum of all mean values. The obtained percentages represent the contribution of each plant or pollinator species to the total flower abundance or number of visits (Table 2).

**Table 2:** Percentage contribution of plant species to the total floral abundance (“FA”) and percentage contribution of pollinator species to the total number of visits (“visits”) for each habitat type explaining at least 80% of the total floral abundance and number of visits, respectively. Indicator species are indicated with a “*”.

| SDHS |  | SNG |  |
| --- | --- | --- | --- |
| Plant species | % FA | Plant species | % FA |
| *Galinsoga quadriradiata* * | 27,5 | *Jacobaea vulgaris* * | 12,8 |
| *Eruca vesicaria* | 12,2 | *Lotus corniculatus* * | 8,7 |
| *Brassica oleracea* | 7,5 | *Trifolium repens* | 8,2 |
| *Fagopyrum esculentum* | 6,9 | *Lythrum salicaria* * | 7,2 |
| *Coriandrum sativum* | 4,5 | *Centaurea jacea* * | 7,1 |
| *Phaseolus vulgaris* | 3,5 | *Epilobium parviflorum* | 5,3 |
| *Trifolium repens* | 3,4 | *Daucus carota* | 4,7 |
| *Trifolium pratense* | 3,2 | *Pulicaria dysenterica* | 3,3 |
| *Origanum vulgare* | 3,0 | *Lysimachia vulgaris* | 3,3 |
| *Mentha* sp. | 2,2 | *Cirsium arvense* * | 3,0 |
| *Apium graveolens* | 2,1 | *Epilobium hirsutum* | 2,8 |
| *Persicaria lapathifolia* | 2,1 | *Symphytum officinale* | 2,8 |
| *Sinapis alba* | 2,0 | *Hypericum perforatum* | 2,6 |
|  |  | *Ranunculus acris* | 2,4 |
|  |  | *Trifolium pratense* | 2,5 |
|  |  | *Hieracium* sp. | 2,2 |
|  |  | *Persicaria maculosa* | 2,2 |
|  | **80,1** |  | **81,1** |
| Pollinator species | % visits | Pollinator species | % visits |
| *Bombus pascuorum* | 34,0 | *Bombus pascuorum* | 30,7 |
| *Eristalis tenax* | 11,7 | *Eristalis tenax* | 8,1 |
| *Bombus lapidarius* | 5,2 | *Bombus lapidarius* | 6,2 |
| *Sphaerophoria scripta* | 5,2 | *Eristalis nemorum* * | 6,1 |
| *Eristalis arbustorum* | 5,1 | *Melitta nigricans* * | 5,8 |
| *Syricta pipiens* * | 4,4 | *Sphaerophoria scripta* | 5,3 |
| *Bombus terrestris* | 4,1 | *Eristalis arbustorum* | 3,5 |
| *Pieris rapae* * | 3,9 | *Myathropa florea* | 3,2 |
| *Hylaeus communis* * | 3,2 | *Bombus terrestris* | 3,0 |
| *Lindenius albilabris* | 2,6 | *Colletes daviesanus* | 2,7 |
| *Eristalis nemorum* | 1,6 | *Helophilus pendulus* * | 2,7 |
|  |  | *Eristalis pertinax* | 1,4 |
|  |  | *Syricta pipiens* | 1,3 |
|  | **81,0** |  | **80,0** |

### Plant-pollinator network

Our network analyses revealed that habitat type significantly affected the degree of specialisation (F = 4.92, p = 0.042; Fig. 4).

**Figure 4:**
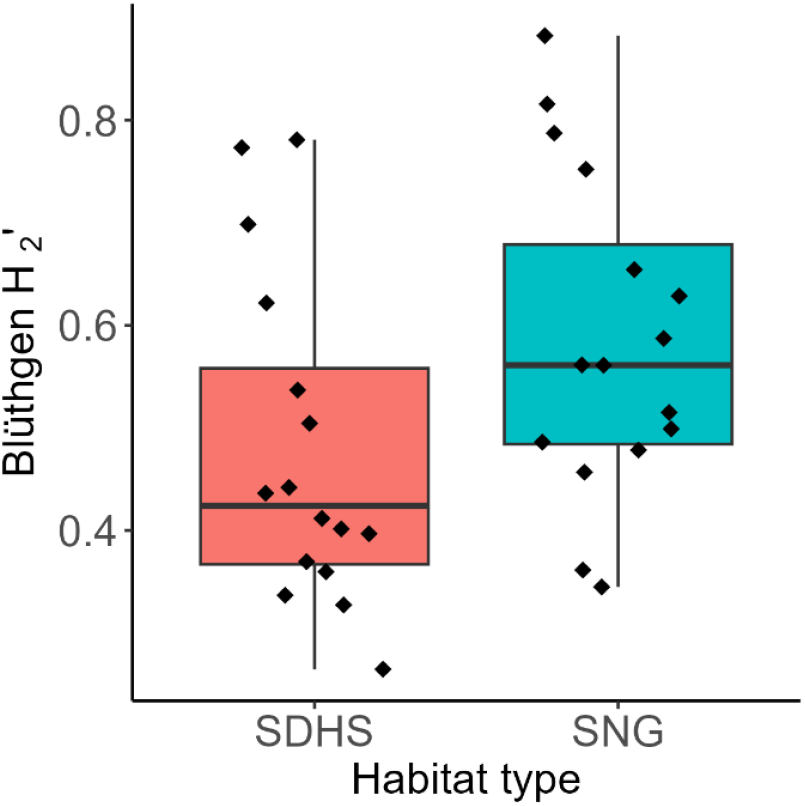
Boxplots of the degree of plant-pollinator network specialisation, expressed as Blüthgen H_2_’ index, for the SDHS (pink) and SNG (blue) habitat type. Each point represents a plant-pollinator network for a combination of sampling location and habitat type.

## Discussion

### General

We found that, compared to semi-natural grasslands (SNG), small-scale, diverse horticultural systems (SDHS) have a similar plant species richness, a higher flower abundance and a distinct floral resource community composition. In addition, we discovered that both pollinator species richness and abundance were similar for both habitat types, yet the composition of the bee and hoverfly community differed between the habitat types. We also identified certain flowering plant and pollinator species as indicator species that were clearly associated with either SDHSs or SNGs. Furthermore, our plant-pollinator network analyses revealed that SNGs harbour a more specialised network compared to SDHSs. We therefore argue that SDHS are a type of agriculture that supports high degrees of floral resources and pollinators in late summer, without taking land out of production or requiring additional management, as is the case for traditional agri-environment schemes. However, SDHSs should be seen as an additional, complementary source of floral resources to those provided by SNG, rather than a substitute, given the importance of these grasslands for pollinators.

### Vegetation

In Flanders, SDHSs provide important floral resources for pollinators in late summer. The floral richness and abundance in these agricultural systems is found to be at least equal to that of SNGs, which are considered to be the most important habitat type for pollinators in late summer. In SNGs (e.g. meadows, permanent pastures), the availability of floral resource is constant during the active period of pollinators. However, their relative importance increases throughout the season, as floral resources in other habitats are generally scarce at that time (Timberlake et al., 2019). Woody semi-natural habitats (SNH), as well as agricultural habitat types like fruit orchards and mass-flowering crops contribute strongly to the provision of floral resource in spring, but their contribution decreases drastically after flowering (Ammann et al., 2024). Besides, landscapes with high amounts of these mass-flowering crops have been shown to negatively affect the reproduction of solitary bees, due to a lack of resources (Eeraerts, Piot, et al., 2021).

To overcome shortages in floral resources, agricultural landscapes can be enriched with flower strips or hedgerows (von Königslöw et al., 2022). Previous research has shown that wildflower strips are as important as extensively managed meadows for pollinator in late summer and that they enhance pollination services in adjacent crops (Feltham et al., 2015; Maurer et al., 2022). Nevertheless, despite scientific evidence and financial compensations, these agri-environmental schemes require a lot of effort, money and space, which makes farmers reluctant to implement them (Batáry et al., 2015; Eeraerts et al., 2020). Therefore, crop diversification has been proposed as a strategy to overcome these challenges by increasing floral resource availability in time and space without taking land out of production (Fijen et al., 2025). All SDHSs we visited implemented at least one diversification measure, such as intercropping, adding trees and shrubs, reducing field sizes, or cultivating new, forgotten or non-crops (e.g. cover crops, green manures). This way, SDHSs provide a diverse set of plant species, which, however, clearly differs from the plant species in SNGs, making SDHSs and SNGs complementary habitats in late summer. Besides, we found that only a limited number of the plant species in each habitat substantially contribute to the total flower abundance, with some species being indicative of a particular habitat type. Conversely, the majority of plant species only contribute marginally to the total flower abundance (Table 2).

The indicator species of SDHS, being *Galinsoga quadriradiata* and *Phacelia tanacetifolia*, together with the number of pollinator visits per plant (Table S1), highlight the importance of non-crops for floral resource provisioning. Cover crops like *Phacelia, Trifolium* sp. and buckwheat may ensure the supply of floral resources for wild pollinators (Bryan et al., 2021; Mallinger et al., 2019), which is confirmed by the great number of pollinator visits to these plants in our study. Weeds may also provide important floral resources, as Balfour & Ratnieks (2022) showed that certain weed species supported a higher abundance and diversity of pollinators compared to wildflower seed mixes or flower strips. Two of the investigated weed species, i.e. *Jacobaea vulgaris* and *Cirsium arvense*, were frequently visited indicator species for SNGs (Table S1). Hence, we suggest that the presence of these and other weed species in agricultural systems could increase floral resource availability. However, further input from practitioners should determine to what extent weeds can be allowed in agricultural systems.

We found that flower abundance in SDHSs was higher compared to SNGs. This could be explained by the presence of certain mass-flowering crops and herbs, such as *Eruca vesicaria, Fagopyrum esculentum, Coriandrum sativum* or flowering *Brassica* sp. However, the number of pollinators in SDHSs did not show a corresponding increase, possibly because the floral resources in this habitat type are less attractive to pollinators than those in SNGs. Grasslands provide the highest amount and diversity of nectar sources (Baude et al., 2016). Data on the attractiveness of the plant species in SDHSs, however, is generally lacking. The FloRes database (Baden-Böhm et al., 2022) provides detailed information on the nectar and pollen production, but it is not exhaustive. In addition, our research exclusively focused on floral resources, with nesting resources not being included. Therefore, to further assess the value of SDHSs for wild pollinating insects, more research should be done on both the attractiveness of floral resources and on the presence of suitable nesting sites.

### Pollinators

In addition to the important floral resource provision in late summer, our findings indicate that SDHSs and SNGs exhibited comparable pollinator species richness and abundance. However, we found that both habitat types differ in their pollinator community composition. Different habitats support distinct sets of wild bee species throughout the season (Mandelik et al., 2012; Maurer et al., 2022). Mallinger et al. (2016) found that bee communities in early-flowering, closed canopy habitats (e.g. woodlands, orchards) were distinguished from those in later-flowering, open habitats (e.g. grasslands, annual croplands). Complementary pollinator communities were also found in sequentially flowering cultivars of sweet cherry (Eeraerts, 2022). However, habitats complement each other not only over time, but also within the same period of time. It has previously been demonstrated that fruit orchards and woody SNHs can complement each other’s floral resources in spring (Eeraerts, Van Den Berge, et al., 2021). Our study adds to this research by showing that agricultural and natural habitats, here represented by SDHSs and SNGs, not only provide distinct sets of plant species, but also serve as habitats for different pollinator species. Additionally, we found that only a limited number of these pollinator species considerably contribute to the total number of visits, with some species being indicative of a particular habitat type. Conversely, the majority of the pollinators were observed to a lesser extent (Table 2).

Indicator species for SDHSs are *Hylaeus communis, Syritta pipiens* and *Pieris rapae*. Of the 80 butterfly individuals we captured, more than half of them were *Pieris rapae*. Together with the other *Pieris* spp. (*P. napi, P. brassicae* and some undefined *Pieris* spp.), this genus represented 80% of the butterfly abundance. Most of these butterflies occurred in the SDHSs, which could explain the significant effect of habitat type on butterfly abundance (Table S4). However, this effect became non-significant after removing outliers (Table S5), which may be mostly driven by the high numbers of *Pieris* spp. compared to other butterfly species. Therefore, we decided to retain the butterfly data solely for combining the results of all pollinators, rather than analysing these data sets separately. This is partly in line with the study by Öckinger & Smith (2007) who excluded these and other butterfly species in order to accurately assess the role of semi-natural grasslands for pollinators in agricultural landscapes.

One of the indicator species for SNGs, *Melitta nigricans*, is a typical oligolectic bee species which exclusively forages on *Lythrum salicaria* (Naturalis, 2024) and was among the most abundant pollinator species in SNGs (Table 2). *Colletes daviesanus*, an oligolectic bee species visiting *Tanacetum vulgare* and a few other *Asteraceae* sp. (Naturalis, 2024), also occurred in high numbers in SNGs. This indicates that SNGs play a crucial role in maintaining populations of rare and specialised pollinator species and was confirmed by the results of the plant-pollinator network analysis. In this analysis we detected that SNGs harbour a more specialised plant-pollinator network compared to that of the SDHSs. This aligns with previous research, showing that the majority of solitary bees encountered in farmlands exhibit a generalist foraging strategy (Wood et al., 2016), while extensively managed meadows support the highest numbers of habitat specialists and rare pollinator species (Maurer et al., 2022). Hadrava et al. (2022) showed that sown flower strips support mostly common generalist bee species, while more specialised solitary bees are found in SNHs. However, in contrast to our visited SDHSs that show a similar species diversity as SNGs, these flower strips were less diverse than SNHs (Hadrava et al., 2022). We therefore assume that crop diversification may be a more effective way of supporting pollinators than sowing flower strips, and also a more economically beneficial measure as no land needs to be taken out of production.

The higher degree of specialisation observed at SNGs may be an overestimation, given that a significant proportion of observations were recorded only once or in a relatively small number. Nevertheless, the same assumption applies to SDHSs. Therefore, we argue that this potential overestimation may be offset and that SDHSs and SNGs are not only complementary in terms of their plant and pollinator community composition, but also with regard to the degree of specialisation of the plant-pollinator networks. Consequently, the combination of both habitat types has the greatest potential to sustain wild pollinators in late summer. This assumption corroborates the findings of Tassoni et al. (2024), who demonstrate that high crop heterogeneity, encompassing both higher field border density and crop diversity at the landscape scale, should be integrated with a sufficient quantity of SNHs in the surrounding landscape in order to encourage and sustain insect populations in agricultural landscapes.

It should be noted that our study was conducted at a single point in time for only one year. To gain a more comprehensive understanding of the value of SDHSs for wild pollinators, this research should be conducted during spring and early summer, and repeated for several years. Furthermore, a comparison of SDHSs with other types of agriculture, such as conventional, biological, or other diversified farming practices, would provide more insight into the relative importance of different agricultural systems. Moreover, only pollinating insects were considered, not the pollination services they deliver, or other organisms delivering other relevant ecosystem services such as pest control. Implementing these objectives in future research, could further evaluate the role of crop diversification strategies for ecosystem services as an alternative for intensive agricultural practices.

## Conclusion

Our results contribute to the limited body of literature on the effects of crop diversification strategies on pollinator conservation in agricultural landscapes. We found that small-scaled, diverse horticultural systems (SDHS) provide a wide range of on-farm floral resources, thereby supporting similar pollinator abundance and richness compared to semi-natural grasslands (SNGs) in late summer. However, the composition of the floral and pollinator community differed between the two habitat types, each characterised by a distinct set of plant and pollinator species. In addition, SNGs were found to support more specialised plant-pollinator networks, confirming the complementary of both habitat types. Our study is therefore one of the first to demonstrate the positive effects of crop diversification as a strategy to enhance agricultural landscapes, while complementing the provision of floral resources of the most important late summer habitat type, i.e. SNGs. Although several other factors determine the intrinsic value of a particular habitat type for biodiversity conservation, such as nesting site suitability for pollinators or the presence of other species like predators for pest control, our research has shown that diversifying agricultural fields can be an effective strategy for maintaining populations wild pollinators in agricultural landscapes.

## Acknowledgement

Maxime Eeraerts was supported as a FWO postdoctoral fellow (grant no. 1210723N).

We thank Augustijn De Ketelaere, Jens D’haeseleer and Jef Hendrix for the determination of wasps, bees and hoverflies, respectively. We also thank Louella Buydens for the help with the network analysis and Jasper Olivier for the help with the field work.

