## Supplemental_Information for "Small-scale, diverse horticultural systems and semi-natural grasslands support complementary pollinator populations"

*Table S1: List of all visited flowering plant species with the number of locations on which they were recorded, if the plant species was visited by pollinators, the numbers of wild pollinators and honeybees per habitat type, and the total number of pollinators recorded on each plant species*

| | | | Wild pollinators | | Honeybees | | |
|---|---|---|---|---|---|---|---|
| **Plant** | **Locations** | **Visited?** | **SDHS** | **SNG** | **SDHS** | **SNG** | **Total** |
| *Achillea millefolium* | 6 | Yes | 0 | 15 | 0 | 0 | 15 |
| *Agastache foeniculum* | 1 | No | 0 | 0 | 0 | 0 | 0 |
| *Agrimonia eupatoria* | 1 | No | 0 | 0 | 0 | 0 | 0 |
| *Allium schoenoprasum* | 2 | Yes | 2 | 0 | 0 | 0 | 2 |
| *Allium* sp. | 1 | No | 0 | 0 | 0 | 0 | 0 |
| *Allium tuberosum* | 2 | Yes | 7 | 0 | 0 | 0 | 7 |
| *Anagallis arvensis* | 1 | No | 0 | 0 | 0 | 0 | 0 |
| *Anchusa officinalis* | 1 | Yes | 0 | 1 | 0 | 0 | 1 |
| *Anethum graveolens* | 1 | Yes | 12 | 0 | 0 | 0 | 12 |
| *Antirrhinum majus* | 1 | No | 0 | 0 | 0 | 0 | 0 |
| *Apium graveolens* | 1 | Yes | 10 | 0 | 0 | 0 | 10 |
| *Arctium lappa* | 1 | Yes | 0 | 2 | 0 | 0 | 2 |
| *Asparagus officinalis* | 1 | No | 0 | 0 | 0 | 0 | 0 |
| *Bellis perennis* | 1 | Yes | 0 | 4 | 0 | 0 | 4 |
| *Borago officinalis* | 3 | Yes | 12 | 0 | 12 | 0 | 24 |
| *Brassica oleracea* | 3 | Yes | 79 | 0 | 24 | 0 | 103 |
| *Brassica* sp. | 1 | Yes | 8 | 0 | 0 | 0 | 8 |
| *Calendula officinalis* | 4 | Yes | 10 | 0 | 1 | 0 | 11 |
| *Callistephus chinensis* | 1 | No | 0 | 0 | 0 | 0 | 0 |
| *Campanula rotundifolia* | 1 | Yes | 0 | 3 | 0 | 0 | 3 |
| *Campanula* sp. | 2 | Yes | 0 | 5 | 0 | 0 | 5 |
| *Capsella bursa-pastoris* | 2 | No | 0 | 0 | 0 | 0 | 0 |
| *Centaurea cyanus* | 3 | Yes | 7 | 0 | 1 | 0 | 8 |
| *Centaurea jacea* | 5 | Yes | 0 | 117 | 0 | 11 | 128 |
| *Centaurea scabiosa* | 1 | No | 0 | 0 | 0 | 0 | 0 |
| *Chenopodium album* | 3 | No | 0 | 0 | 0 | 0 | 0 |
| *Cichorium intybus* | 3 | Yes | 1 | 0 | 0 | 0 | 1 |
| *Cirsium arvense* | 9 | Yes | 10 | 51 | 0 | 6 | 67 |
| *Cirsium palustre* | 3 | Yes | 0 | 33 | 0 | 0 | 33 |
| *Cirsium vulgare* | 1 | Yes | 0 | 1 | 0 | 0 | 1 |

| | | | | | | | |
|---|---|---|---|---|---|---|---|
| *Consolida regalis* | 1 | No | 0 | 0 | 0 | 0 | 0 |
| *Convolvulus arvensis* | 5 | Yes | 0 | 1 | 0 | 1 | 2 |
| *Convolvulus sepium* | 2 | Yes | 0 | 1 | 0 | 0 | 1 |
| *Convolvulus* sp. | 1 | No | 0 | 0 | 0 | 0 | 0 |
| *Coriandrum sativum* | 5 | Yes | 50 | 0 | 3 | 0 | 53 |
| *Cosmos bipinnatus* | 1 | Yes | 2 | 0 | 0 | 0 | 2 |
| *Crepis capillaris* | 7 | Yes | 4 | 6 | 0 | 0 | 10 |
| *Cucurbita* sp. | 3 | Yes | 5 | 0 | 15 | 0 | 20 |
| *Cynara cardunculus* | 1 | Yes | 0 | 0 | 5 | 0 | 5 |
| *Daucus carota* | 6 | Yes | 2 | 50 | 0 | 0 | 52 |
| *Dianthus armeria* | 1 | No | 0 | 0 | 0 | 0 | 0 |
| *Dianthus deltoides* | 1 | Yes | 0 | 6 | 0 | 0 | 6 |
| *Dipsacus fullonum* | 1 | Yes | 0 | 2 | 0 | 0 | 2 |
| *Echinochloa crus-galli* | 1 | No | 0 | 0 | 0 | 0 | 0 |
| *Epilobium hirsutum* | 3 | Yes | 0 | 10 | 0 | 1 | 11 |
| *Epilobium parviflorum* | 6 | Yes | 0 | 9 | 1 | 1 | 11 |
| *Erigeron canadensis* | 7 | No | 0 | 0 | 0 | 0 | 0 |
| *Eruca vesicaria* | 3 | Yes | 89 | 0 | 17 | 0 | 106 |
| *Eupatorium cannabinum* | 3 | Yes | 0 | 9 | 0 | 7 | 16 |
| *Fagopyrum esculentum* | 3 | Yes | 43 | 0 | 4 | 0 | 47 |
| *Foeniculum vulgare* | 5 | Yes | 21 | 0 | 0 | 0 | 21 |
| *Galinsoga parviflora* | 1 | Yes | 1 | 0 | 0 | 0 | 1 |
| *Galinsoga quadriradiata* | 11 | Yes | 28 | 0 | 22 | 0 | 50 |
| *Galium verum* | 1 | No | 0 | 0 | 0 | 0 | 0 |
| *Geranium molle* | 1 | No | 0 | 0 | 0 | 0 | 0 |
| *Geranium* sp. | 2 | Yes | 0 | 1 | 0 | 0 | 1 |
| *Glebionis segetum* | 1 | Yes | 12 | 0 | 0 | 0 | 12 |
| *Gnaphalium uliginosum* | 1 | No | 0 | 0 | 0 | 0 | 0 |
| *Helianthus annuus* | 3 | Yes | 40 | 0 | 7 | 0 | 47 |
| *Heracleum sphondylium* | 8 | Yes | 0 | 80 | 0 | 6 | 86 |
| *Hieracium* sp. | 3 | Yes | 0 | 8 | 0 | 22 | 30 |
| *Hypericum perforatum* | 4 | Yes | 0 | 11 | 1 | 0 | 12 |
| *Hypochaeris radicata* | 2 | Yes | 0 | 3 | 0 | 0 | 3 |
| *Hyssopus officinalis* | 2 | Yes | 10 | 0 | 0 | 0 | 10 |
| *Jacobaea vulgaris* | 6 | Yes | 0 | 63 | 0 | 1 | 64 |
| *Knautia arvensis* | 1 | Yes | 0 | 3 | 0 | 0 | 3 |
| *Lactuca serriola* | 1 | No | 0 | 0 | 0 | 0 | 0 |
| *Lamium album* | 1 | No | 0 | 0 | 0 | 0 | 0 |
| *Lamium purpureum* | 8 | Yes | 10 | 0 | 0 | 0 | 10 |
| *Lavendula* sp. | 1 | Yes | 1 | 0 | 0 | 0 | 1 |
| *Leontodon* sp. | 4 | Yes | 8 | 1 | 0 | 0 | 9 |
| *Leonurus cardiaca* | 1 | Yes | 13 | 0 | 4 | 0 | 17 |
| *Leucanthemum vulgare* | 3 | Yes | 0 | 8 | 0 | 0 | 8 |
| *Levisticum officinale* | 1 | Yes | 1 | 0 | 0 | 0 | 1 |
| *Linaria vulgaris* | 1 | No | 0 | 0 | 0 | 0 | 0 |
| *Lotus corniculatus* | 10 | Yes | 3 | 36 | 0 | 0 | 39 |
| *Lycopus europaeus* | 4 | Yes | 0 | 1 | 0 | 6 | 7 |
| *Lysimachia vulgaris* | 2 | Yes | 0 | 8 | 0 | 0 | 8 |

| | | | | | | | |
|---|---|---|---|---|---|---|---|
| *Lythrum salicaria* | 9 | Yes | 0 | 125 | 0 | 3 | 128 |
| *Malva sylvestris* | 4 | Yes | 17 | 0 | 6 | 0 | 23 |
| *Matricaria chamomilla* | 2 | No | 0 | 0 | 0 | 0 | 0 |
| *Matricaria* sp. | 4 | No | 0 | 0 | 0 | 0 | 0 |
| *Medicago lupulina* | 1 | No | 0 | 0 | 0 | 0 | 0 |
| *Medicago sativa* | 2 | Yes | 25 | 0 | 0 | 0 | 25 |
| *Mentha aquatica* | 2 | Yes | 19 | 8 | 7 | 4 | 38 |
| *Mentha* sp. | 2 | Yes | 16 | 0 | 8 | 0 | 24 |
| *Mentha spicata* | 3 | Yes | 2 | 0 | 0 | 0 | 2 |
| *Mentha x piperita* | 2 | No | 0 | 0 | 0 | 0 | 0 |
| *Mentha x rotundifolia* | 1 | Yes | 18 | 0 | 1 | 0 | 19 |
| *Myosotis* sp. | 1 | No | 0 | 0 | 0 | 0 | 0 |
| *Myosoton aquaticum* | 1 | No | 0 | 0 | 0 | 0 | 0 |
| *Nigella damascena* | 1 | Yes | 0 | 0 | 1 | 0 | 1 |
| *Orchis* sp. | 1 | No | 0 | 0 | 0 | 0 | 0 |
| *Origanum majorana* | 1 | Yes | 2 | 0 | 0 | 0 | 2 |
| *Origanum vulgare* | 5 | Yes | 43 | 21 | 5 | 0 | 69 |
| *Oxalis corniculata* | 1 | No | 0 | 0 | 0 | 0 | 0 |
| *Papaver rhoeas* | 2 | No | 0 | 0 | 0 | 0 | 0 |
| *Persicaria lapathifolia* | 3 | Yes | 2 | 0 | 1 | 0 | 3 |
| *Persicaria maculosa* | 9 | Yes | 10 | 3 | 0 | 0 | 13 |
| *Phacelia tanacetifolia* | 5 | Yes | 30 | 0 | 4 | 0 | 34 |
| *Phalaris arundinacea* | 1 | No | 0 | 0 | 0 | 0 | 0 |
| *Phaseolus vulgaris* | 3 | Yes | 33 | 0 | 3 | 0 | 36 |
| *Physalis* sp. | 1 | Yes | 15 | 0 | 0 | 0 | 15 |
| *Pimpinella major* | 1 | Yes | 0 | 14 | 0 | 0 | 14 |
| Plant 1 | 1 | Yes | 1 | 0 | 0 | 0 | 1 |
| Plant 4 | 1 | No | 0 | 0 | 0 | 0 | 0 |
| *Plantago lanceolata* | 9 | Yes | 0 | 4 | 0 | 0 | 4 |
| *Poaceae* sp. | 6 | No | 0 | 0 | 0 | 0 | 0 |
| *Prunella vulgaris* | 3 | No | 0 | 0 | 0 | 0 | 0 |
| *Pulicaria dysenterica* | 4 | Yes | 2 | 22 | 0 | 5 | 29 |
| *Ranunculus acris* | 6 | Yes | 1 | 14 | 0 | 0 | 15 |
| *Ranunculus flammula* | 1 | No | 0 | 0 | 0 | 0 | 0 |
| *Ranunculus repens* | 2 | No | 0 | 0 | 0 | 0 | 0 |
| *Ranunculus* sp. | 2 | No | 0 | 0 | 0 | 0 | 0 |
| *Raphanus sativus* | 2 | Yes | 24 | 0 | 2 | 0 | 26 |
| *Rorippa palustris* | 2 | No | 0 | 0 | 0 | 0 | 0 |
| *Rubus fruticosus* | 1 | No | 0 | 0 | 0 | 0 | 0 |
| *Rubus idaeus* | 1 | Yes | 23 | 0 | 5 | 0 | 28 |
| *Salvia nemorosa* | 1 | Yes | 6 | 0 | 0 | 0 | 6 |
| *Satureja hortensis* | 2 | Yes | 18 | 0 | 0 | 0 | 18 |
| *Scabiosa columbaria* | 1 | Yes | 0 | 35 | 0 | 0 | 35 |
| *Senecio vulgaris* | 5 | Yes | 1 | 0 | 0 | 0 | 1 |
| *Silene latifolia* | 1 | No | 0 | 0 | 0 | 0 | 0 |
| *Sinapis alba* | 1 | Yes | 18 | 0 | 1 | 0 | 19 |
| *Solanum lycopersicum* | 1 | Yes | 1 | 0 | 0 | 0 | 1 |
| *Solanum melongena* | 1 | Yes | 5 | 0 | 0 | 0 | 5 |

| | | | | | | | |
|---|---|---|---|---|---|---|---|
| *Solanum nigrum* | 7 | Yes | 7 | 1 | 0 | 0 | 8 |
| *Sonchus arvensis* | 1 | Yes | 31 | 0 | 1 | 0 | 32 |
| *Sonchus asper* | 1 | No | 0 | 0 | 0 | 0 | 0 |
| *Sonchus oleraceus* | 2 | No | 0 | 0 | 0 | 0 | 0 |
| *Sonchus* sp. | 6 | Yes | 1 | 0 | 0 | 0 | 1 |
| *Stachys palustris* | 1 | Yes | 0 | 3 | 0 | 0 | 3 |
| *Stellaria aquatica* | 1 | Yes | 0 | 6 | 0 | 1 | 7 |
| *Stellaria graminea* | 1 | No | 0 | 0 | 0 | 0 | 0 |
| *Stellaria holostea* | 1 | No | 0 | 0 | 0 | 0 | 0 |
| *Stellaria media* | 5 | Yes | 0 | 1 | 0 | 0 | 1 |
| *Stellaria* sp. | 1 | No | 0 | 0 | 0 | 0 | 0 |
| *Succisa pratensis* | 1 | No | 0 | 0 | 0 | 0 | 0 |
| *Symphytum officinale* | 8 | Yes | 10 | 37 | 0 | 0 | 47 |
| *Tanacetum parthenium* | 1 | Yes | 5 | 0 | 0 | 0 | 5 |
| *Tanacetum vulgare* | 3 | Yes | 0 | 32 | 0 | 0 | 32 |
| *Taraxacum officinale* | 5 | Yes | 12 | 0 | 0 | 0 | 12 |
| *Thymus vulgaris* | 4 | Yes | 1 | 0 | 2 | 0 | 3 |
| *Trifolium dubium* | 1 | No | 0 | 0 | 0 | 0 | 0 |
| *Trifolium pratense* | 9 | Yes | 28 | 37 | 1 | 0 | 66 |
| *Trifolium repens* | 9 | Yes | 17 | 30 | 2 | 3 | 52 |
| *Tropaeolum majus* | 1 | No | 0 | 0 | 0 | 0 | 0 |
| *Urtica dioica* | 1 | No | 0 | 0 | 0 | 0 | 0 |
| *Urtica* sp. | 4 | No | 0 | 0 | 0 | 0 | 0 |
| *Veronica persica* | 1 | No | 0 | 0 | 0 | 0 | 0 |
| *Vicia cracca* | 1 | Yes | 0 | 6 | 0 | 0 | 6 |
| *Viola arvensis* | 1 | No | 0 | 0 | 0 | 0 | 0 |
| *Viola* sp. | 2 | No | 0 | 0 | 0 | 0 | 0 |
| *Viola tricolor* | 1 | No | 0 | 0 | 0 | 0 | 0 |
| | | | **945** | **948** | **167** | **78** | **2138** |

*Table S2: Overview of all sampled landscapes and the corresponding sampling date. "Start of sampling" refers to the start time of the pollinator survey. "Mean temperature" is the mean value of the four temperatures measured during transect walks at each transect*

| Location | Sampling date | Start of sampling | Mean temperature |
|---|---|---|---|
| 1 | 25/07/2023 | 14:30 | 19°C |
| 2 | 26/07/2023 | 11:10 | 18.7°C |
| 3 | 28/07/2023 | 15:00 | 21°C |
| 4 | 29/07/2023 | 14:30 | 22.5°C |
| 1 | 30/07/2023 | 13:50 | 20.3°C |
| 5 | 01/08/2023 | 12:00 | 19°C |
| 6 | 09/08/2023 | 13:50 | 20.5°C |
| 7 | 10/08/2023 | 13:00 | 22.5°C |
| 8 | 11/08/2023 | 13:00 | 17.5°C |
| 9 | 14/08/2023 | 11:30 | 24.3°C |
| 10 | 15/08/2023 | 10:40 | 23.3°C |
| 11 | 16/08/2023 | 11:10 | 22.3°C |
| 12 | 18/08/2023 | 11:00 | 21.3°C |
| 13 | 24/08/2023 | 10:30 | 22.7°C |
| 14 | 29/08/2023 | 11:50 | 18°C |
| 15 | 04/09/2023 | 11:40 | 22°C |
| 16 | 05/09/2023 | 12:10 | 27°C |

*Table S3: Mean and SE for each response variable: species richness ("SR") and flower abundance ("FA") of the visited flowering vegetation, and species richness and abundance ("AB") of all pollinators, bees and hoverflies.*

| Response | Habitat type | Mean | SE |
|---|---|---|---|
| SR Vegetation | SDHS | 7.56 | 0.62 |
| | SNG | 7.63 | 0.60 |
| FA Vegetation | SDHS | 2505.19 | 448.39 |
| | SNG | 820.56 | 121.05 |
| SR All | SDHS | 14.44 | 1.04 |
| | SNG | 15.25 | 1.42 |
| SR Bee | SDHS | 5.25 | 0.64 |
| | SNG | 6.06 | 0.65 |
| SR Hoverfly | SDHS | 5.75 | 0.78 |
| | SNG | 5.81 | 0.70 |
| AB All | SDHS | 59.06 | 5.22 |
| | SNG | 59.25 | 4.58 |
| AB Bee | SDHS | 31.25 | 3.10 |
| | SNG | 33.63 | 4.61 |
| AB Hoverfly | SDHS | 20.38 | 4.73 |
| | SNG | 20.75 | 3.31 |

*Table S4: Summary of the results of the generalized linear mixed-effect models for the effect of the habitat type on pollinator abundance (“AB”) and species richness (“SR”) for butterflies and wasps. “Response” and “Fixed are the response variable and fixed variable of the model, respectively. The reported statistic are the z-value (“z”) and the corresponding p-value (“p”), together with the mean (“Mean”) and standard error (“SE”) for each habitat type.*

| Response | Fixed | z | p | Habitat type | Mean | SE |
|---|---|---|---|---|---|---|
| AB butterfly | Habitat type | -2.32 | 0.02 | CSA | 3.85 | 1.67 |
| | | | | SNG | 2.14 | 0.44 |
| AB wasp | Habitat type | -2.12 | 0.03 | CSA | 5.31 | 1.59 |
| | | | | SNG | 4.36 | 0.75 |
| SR butterfly | Habitat type | 0.57 | 0.57 | CSA | 1.31 | 0.13 |
| | | | | SNG | 1.57 | 0.25 |
| SR wasp | Habitat type | -0.11 | 0.91 | CSA | 2.92 | 0.61 |
| | | | | SNG | 2.91 | 0.56 |

*Table S5: Summary of the results of the generalized linear-mixed effect models for the effect of habitat type on the species richness ("SR") and abundance ("AB") of the different pollinator types. “Response” and “Fixed” are the response variable and fixed variable of the model, respectively. The reported statistics are the z-value (“z”) with its corresponding p-value (“p”).*

| Response | Fixed | z | p |
|---|---|---|---|
| AB all | Habitat type | -1.57 | 0.12 |
| AB bee | Habitat type | -0.27 | 0.79 |
| AB butterfly | Habitat type | 0.33 | 0.74 |
| AB wasp | Habitat type | -17.21 | < 0.001 |
| SR all | Habitat type | -0.58 | 0.56 |
| SR bee | Habitat type | 0.80 | 0.42 |
| SR hoverfly | Habitat type | -0.57 | 0.57 |
| SR butterfly | Habitat type | 0.17 | 0.87 |
| SR wasp | Habitat type | 0.71 | 0.48 |
